# Cis-regulatory variation and transcription factor binding contribute to allelic genotype-by-environment interactions for gene expression in maize

**DOI:** 10.64898/2026.07.31.742149

**Authors:** Sontosh K. Deb, Ty Thomas, Jordan Cummings, Katelyn Rumley, Melissa A. Draves, James Holland, Jacob D. Washburn, Sherry Flint-Garcia, Joseph L. Gage

**Affiliations:** Department of Crop and Soil Sciences, North Carolina State University, Raleigh, NC, USA; NC Plant Sciences Initiative, North Carolina State University, Raleigh, NC, USA; Division of Plant Science and Technology, University of Missouri, Columbia, MO, USA; USDA-ARS Plant Science Research Unit, Raleigh, NC, USA; USDA-ARS Plant Genetics Research Unit, Columbia, MO, USA

**Keywords:** genotype by environment, GxE, allele by environment, AxE, allele specific expression, ASE, maize, cis-regulatory variation

## Abstract

Genotype-by-environment interactions (GxE), or differences in how genotypes perform across varying environments, are a pervasive source of phenotypic variation and underlie differences in local adaptation. Though GxE is well characterized across kingdoms of life, less is known about what causes GxE interactions, particularly at the molecular level. In this study, we use allele-specific gene expression estimates in a maize (*Zea mays* L.) B73 x Mo17 hybrid to isolate cis-regulatory effects on gene expression for each of the two parental alleles. The hybrid was grown in two environments, and expression differences between the parental alleles were used to characterize allele-by-environment (AxE) interactions and study the influence of gene-proximal sequence variation on transcript abundance AxE. We tested the hypothesis that gene-proximal sequence variation can cause GxE in gene expression by modifying transcription factor binding. Our results show that sequence variation in gene promoter regions has a small but consistent enrichment in genes that show transcriptional AxE. Further, we demonstrate that differential transcription factor binding potential caused by sequence variation is also enriched in AxE genes. Predictive models trained on sequence and transcription factor binding variation show that while these features contain some information about whether a gene will show transcriptional AxE, they alone are not sufficient to reliably distinguish AxE genes. These findings support the hypothesis that gene expression GxE can be caused by sequence variation that modifies transcription factor binding, while also reinforcing the complex and context-specific nature of GxE interactions.

## Introduction

Understanding how a specific plant genotype performs in a given environment is critical for plant breeding, assisted migration, and predicting how crops will adapt to novel conditions(van Kleunen and Fischer 2005; Velotta and Cheviron 2018; Teressa et al. 2021). Genotype-by-environment interactions (G×E), where genetically distinct individuals show different phenotypic responses to environmental change, are a source of variation that can influence the environment-specific phenotype of an individual. Maize (*Zea mays* L.) is well-suited for studying GxE because of its agricultural significance, well developed genomic resources, and because inbred and hybrid genotypes can be easily replicated in multi-environment field trials(Kusmec et al. 2018). In a previous study of 22 traits in field-grown maize inbred lines, GxE accounted for an average of 15% and a maximum of 69% of per-plot variation(Hung et al. 2012).

Despite the importance of GxE and its extensive characterization in quantitative traits such as fitness or flowering time(Rogers et al. 2021; Mora-Poblete et al. 2023; Napier et al. 2023; Siddiq et al. 2024), our understanding of GxE interactions at the molecular level remains largely limited to model organisms(Smith and Kruglyak 2008; Grishkevich et al. 2012; Huang et al. 2020; Yun et al. 2025). Gene expression, protein abundance, and other molecular traits ultimately drive macroscopic phenotypes(Jakobson and Jarosz 2019), meaning molecular GxE may underpin organismal-level responses. Indeed, some studies show genes exhibiting a transcriptional response to the environment are up to seven times more likely to be associated with complex phenotypes(Moyerbrailean et al. 2016; Boye et al. 2024), highlighting why deciphering molecular GxE is necessary to improve whole-plant traits.

The regulatory mechanisms underlying transcriptional GxE remain difficult to generalize(Lovell et al. 2016; Albert et al. 2018; Ballinger et al. 2023; Mack et al. 2025). Environment-dependent expression differences can arise through both local variation near the affected gene and distal factors, such as transcription factors or other diffusible regulatory components(Wittkopp and Kalay 2011; Signor and Nuzhdin 2018; Mack et al. 2025). Though local/distal and cis/trans are sometimes used interchangeably, we use local/distal to categorize regulatory variation near/far from focal genes, and refer to cis/trans when mechanistic connection is known, since cis- and trans-regulation can occur locally or distally(Rockman and Kruglyak 2006).

Evidence from different systems suggests that the relative contribution of both local and distal mechanisms is context-dependent. In maize, variation in gene expression responses to cold and heat stress among inbred lines was explained by both local and distal regulatory effects, but local regulatory variation was more common for both steady-state and stress-responsive expression differences(Waters et al. 2017). Similarly, drought-response studies in Arabidopsis(Cubillos et al. 2014) and switchgrass(Lovell et al. 2016) found a bias toward local regulatory variation, both when comparing accessions and when comparing treatment and control plants. In contrast, several studies have found that distal regulatory effects are more sensitive to environmental conditions and often explain a larger fraction of expression plasticity or expression GxE(Albert et al. 2018; Ballinger et al. 2023; Mack et al. 2025). Together, these studies suggest that although distal regulatory effects can make major contributions to environment-dependent transcriptional responses(Smith and Kruglyak 2008), local regulatory effects are repeatedly detected and may represent an important component of transcriptional GxE.

Several approaches have been used to distinguish local and distal regulatory contributions to gene expression variation. Expression quantitative trait locus (eQTL) mapping treats transcript abundance as a quantitative trait and can identify genomic regions associated with variation in expression levels, including local and distal eQTLs(Schadt et al. 2003; Nica and Dermitzakis 2013). Response-eQTL approaches extend this framework to environment-dependent expression by identifying genetic variants whose effects on expression change under specific environmental or cellular conditions(Barreiro et al. 2012; Fairfax et al. 2014; Lee et al. 2014; Moyerbrailean et al. 2016). However, eQTL-based approaches often require large sample sizes and can have limited resolution for distinguishing whether a local eQTL reflects a true cis-regulatory variant or a nearby trans-acting regulator(Rockman and Kruglyak 2006; Moyerbrailean et al. 2016).

Allele-specific expression (ASE) provides a complementary framework for resolving these problems by comparing expression differences between parental alleles in F1 hybrids(Wittkopp et al. 2004; Landry et al. 2007; Castel et al. 2015). Because both parental alleles are measured within the same hybrid individual and are exposed to a shared cellular and trans-regulatory environment, biased expression between alleles in the F1 is interpreted as evidence for cis-regulatory, rather than simply local, divergence(Cowles et al. 2002; Yan et al. 2002; Wittkopp et al. 2004). ASE has been applied across several plant systems to study regulatory variation in gene expression(Springer and Stupar 2007; Cubillos et al. 2014; Lovell et al. 2016; Waters et al. 2017; Albert et al. 2018; Shao et al. 2019; Hu et al. 2022).

Building on this framework, we leverage ASE to estimate genetic and GxE effects on gene expression of the two parental alleles in a single maize hybrid, B73xMo17. Using ASE to estimate gene expression of both parental alleles provides two contrasting allelic effects (B73 and Mo17) within a single plant, minimizing noise due to developmental differences, tissue composition, micro-environmental variation, and trans-regulatory effects that would be present if we studied the two parents as inbred lines. Our study uses ASE across two environmental conditions to identify genes whose allelic responses vary by environment, providing evidence for cis-regulatory contributions to transcriptional GxE. Because in this experiment we are studying interaction effects using two alleles within a single genotype (the B73xMo17 hybrid), we will refer to interaction effects from this particular experiment as allele-by-environment interactions (AxE), which we consider to be a type of GxE. We test whether the transcriptional AxE identified in this study is generalizable to plant-level phenotypic GxE in a biparental population of inbred lines from the same two parents: the intermated B73xMo17 population (IBM). Finally, we use reference-quality genome assemblies for the parental inbreds and publicly available DNA Affinity Purification sequencing (DAP-seq) data (Galli, Chen, et al. 2025) to identify and model sequence variation and transcription factor binding sites associated with transcriptional AxE.

## Results & Discussion

### Allele-specific expression reveals patterns of cis-regulated transcriptional AxE

The goal for this study was to analyze cis-regulatory determinants of transcript abundance AxE. Sampling F1 hybrids between B73 and Mo17 and estimating ASE provides two convenient advantages. First, transcript abundance estimates for each parental allele are derived from the same population of cells in sampled tissue, removing any variation due to plot-to-plot microenvironment, developmental differences, or tissue heterogeneity that might be seen when sampling genetically distinct inbreds. Second, using F1 hybrids minimizes trans-regulated variation, allowing us to focus on linking cis-regulatory variation to transcript abundance.

Initial modeling of allele-specific counts as a function of allele, environment, and allele x environment (AxE) effects resulted in p-value distributions for the AxE term that were skewed toward 1, violating the assumption of uniformly distributed p-values and indicating that the model was improperly specified, perhaps due to dependence between alleles from the same sample. We adopted the modeling strategy described in(Love 2017), which models AxE interactions based on the ratio of allelic counts within each sample. This modeling framework means that the statistical AxE test is for changes in the ratio of parental alleles between environments, meaning that certain change-in-magnitude reaction norms where the alternate and reference alleles have the same ratio but different mean expression levels are not identified as AxE interactions. For example, if the count of Mo17:B73 reads for a hypothetical gene is 20:10 in one environment and 200:100 in another, that gene would not be identified as having significant AxE under this allelic ratio model, although it may have a significant AxE under other modelling strategies. The change-in-ratio model resulted in a distribution of AxE p-values that was largely uniform, with an enrichment of low p-value genes that we interpret to represent genes with AxE effects.

Our allelic ratio model identified 219 genes (1.6% of tested genes) showing significant AxE effects for transcript abundance (FDR=0.1; Figure 1A; Supplementary File 1). Due to differences in developmental timing between the two locations (NC sampled 11 days and 166 growing degree units later than MO), we cannot confidently attribute all interaction effects to environmental influences; there may be some degree of allele-by-development interactions entangled with our AxE signal. However, the identified ‘AxE genes’ still represent context-specific changes in allelic gene regulation.

**Figure 1:**
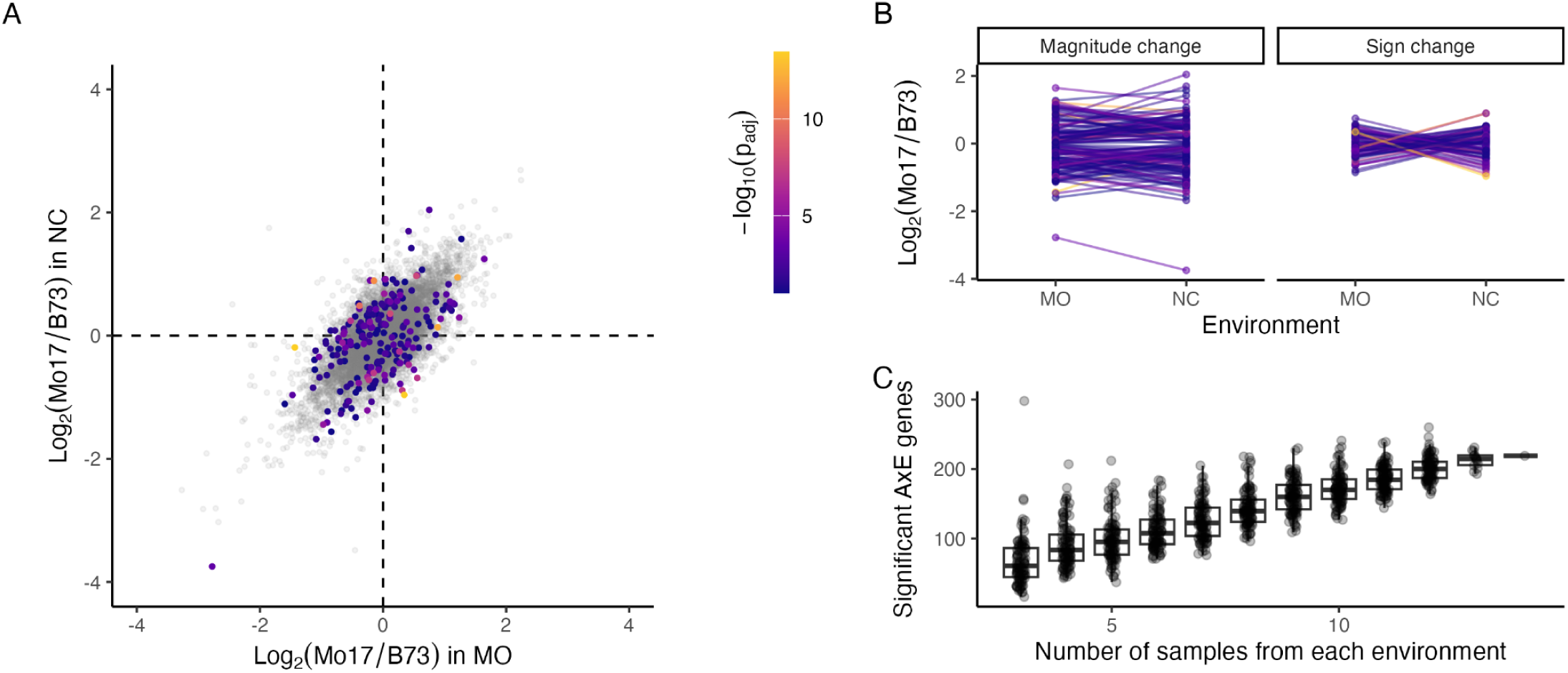
A) Scatterplot showing allele-specific transcript abundance ratios in Missouri versus North Carolina. Genes with significant allele-environment interactions (AxE) (FDR=0.1) are colored according to their Benjamini-Hochberg adjusted p-value. Thirteen non-significant points outside the range of the axes were excluded from the plot. B) Change in allele-specific transcript abundance ratio between environments for 219 significant AxE genes. Genes are split between panels based on whether the sign of their log-ratio changes between environments. C) Power to detect significant AxE interactions increases linearly with sample size. Sample sizes between n=3 and n=14 (full dataset) were used to test for AxE interactions for transcript abundance in B73xMo17 hybrids grown in replicated field trials in North Carolina and Missouri.

Because our allele-specific experimental design limits distal effects, which have been previously shown to comprise the majority of GxE eQTL in some model species (Li et al. 2006; Smith and Kruglyak 2008; Liu et al. 2020), we expected to identify a limited number of AxE genes in this study. However, the proportion of AxE genes we identified is still similar to those reported in previous experiments in maize (0.05%)(Chen et al. 2015), Brachypodium (2%)(Yun et al. 2025), Drosophila (2-3%)(Huang et al. 2020), and yeast (4-40%)(Landry et al. 2006; Smith and Kruglyak 2008), all of which include both proximal and distal effects. High replication (n=14 per environment) and differences in statistical approaches may account for the unexpectedly high number of cis-regulated AxE genes we identified.

Of those 219 AxE genes, 77 (35%) showed changes in which parental allele was more highly expressed, whereas for 142 (65%), the same allele had higher expression in both environments, but the magnitude of the allelic ratio changed (Figure 1B). This contrasts with a study of *Drosophila melanogaster* inbred lines, in which almost no genes showed sign change between environments(Huang et al. 2020). Notably, genes showing a switch in the higher expressed allele generally had greater absolute fold-changes than genes that showed changes in magnitude (Mann-Whitney p=4.7e-5).

### Power analysis of transcriptional GxE

To inform future studies of GxE and/or AxE effects in gene expression, we evaluated the effect of sample size on power to identify statistically significant AxE effects. Because we had 14 samples of B73xMo17 in each environment, we were able to downsample our data to simulate a dataset with fewer replicates. In addition to providing information that may be useful for designing future experiments, we also wanted to answer the question: is there a finite ‘set’ of AxE genes that can be detected when contrasting a particular pair of alleles in a particular set of environments, or is there a very large number of AxE genes whose detection is more dependent on sample size and power to detect increasingly small effects? If there is a fixed number of AxE genes, we’d expect the number of significant genes to saturate at some point as sample size increases. On the other hand, if AxE interactions are not limited to a relatively small set of differentially-responsive genes, we would expect the number of significant AxE genes to continually increase as we increase our sample size.

In this particular experiment, as we increased our sample size from three to 14 replicates per location, the number of AxE genes detected as significant rose linearly from 60 to 219 significant genes (FDR=0.1; representing 0.5% - 1.6% of tested genes) and did not show any sign of leveling off (Figure 1C; Supplementary File 2). This means that either AxE interactions are pervasive but of varying effect size, with smaller effects being increasingly more difficult to detect; or it means that we are still underpowered at 14 replicates to identify the full fixed set of AxE genes. From a practical standpoint, even if a fixed set of AxE genes does exist, it is unlikely that many future studies will have sufficient sample size (n>14) to find them all. Though these specific results are a product of the particular genotypes being compared and then environments included, future studies will need to weigh the tradeoff between replication cost and power to detect cis-regulated GxE, and can use the findings presented here as prior information for making those experimental design decisions.

### Transcriptional AxE interactions do not co-localize with phenotypic GxE from independent experiments

We wanted to test whether the observed AxE patterns for transcript abundance could be statistically connected to GxE for plant-level phenotypes. Using phenotypic data from multi-environment trials of the intermated B73xMo17 (IBM) population, a set of recombinant inbred lines (RILs) derived from the same parents as the hybrids in this study (Lee et al. 2002), we mapped quantitative trait loci (QTL) for GxE interaction effects for 19 traits in up to 10 environments (Hung et al. 2012). We found no significant overlap between AxE genes from our transcriptomic study and recombination bins with significant whole-plant phenotype GxE QTL.

It is possible that testing for transcriptomic AxE in genetically uniform B73xMo17 hybrids, where both alleles share a trans-regulatory background, inherently reveals a different set of genes (ie, preferentially cis-regulated) than phenotyping trials using recombinant inbred lines, which segregate for combinations of cis-regulatory, trans-regulatory and polygenic background effects. Previous experiments mapping transcriptomic GxE and plasticity in biparental populations have identified a large influence from trans effects(Li et al. 2006; Smith and Kruglyak 2008; Lensink et al. 2025). If a preponderance of trans-regulatory QTL is also the case for plant-level phenotypes, then that would explain the lack of overlap between GxE trait QTL and the cis-regulated AxE genes that are the focus of this experiment. The distinction in regulatory architecture is particularly notable in the context of maize breeding, where families of inbred lines are evaluated and compared to each other per se or as testcrosses across multiple locations, but with the ultimate goal of releasing hybrids that must perform reliably in a target range of environments.

### AxE genes are enriched for gene-proximal sequence variation

One hypothesis for how sequence variation drives transcriptional GxE is that differences in proximal regulatory sequences cause different patterns of transcriptional regulation under different contexts(Boye et al. 2024). Under this hypothesis, we expect that genes showing transcriptional AxE would be enriched for sequence variation that causes differential TF binding both between alleles and between conditions. To begin testing whether and how proximal sequence variation influences GxE for gene expression, we tested if our 219 genes with significant AxE effects were enriched for sequence variation in their promoters. We tested for enrichment of SNPs, small indels (<50bp), or large indels (≥50bp) within 500bp of the transcription start site (TSS) in AxE genes, compared to a background set of genes.

We found mild enrichment for the presence of small and large indels in AxE gene promoters relative to background genes and no evidence that AxE genes have SNPs more frequently than background (Figure 2; Supplementary File 3). However, when considering the number of variants in promoter regions, rather than just presence/absence, we found all variant types are more frequent in AxE gene promoters relative to background (Mann-Whitney p=0.02 for large indels, 2e-6 for small indels, and 6e-5 for SNPs; Figure 2). These results differ from findings in *C. elegans* that show an enrichment of SNPs in the promoter of genes with expression variation across genotypes but not among genes with GxE interactions, putatively due to predominant distal regulation of GxE genes(Grishkevich et al. 2012). Our results may differ because we are focused on cis-regulated AxE genes. Other studies have suggested that GxE genes are highly regulated(Promislow 2005; Landry et al. 2006; Grishkevich et al. 2012), but it is also possible that AxE genes we identified are under less constraint than background genes, as previously shown for Arabidopsis genes with condition-specific expression(Roberts and Josephs 2023), and thus have accumulated more mutations. The enrichment of promoter sequence variants indicates a potential mechanism for observed transcriptional GxE. To test this, we next used publicly available DAP-seq data to test whether AxE expression patterns may be driven by differential transcription factor (TF) binding where sequences differ between the two alleles.

**Figure 2:**
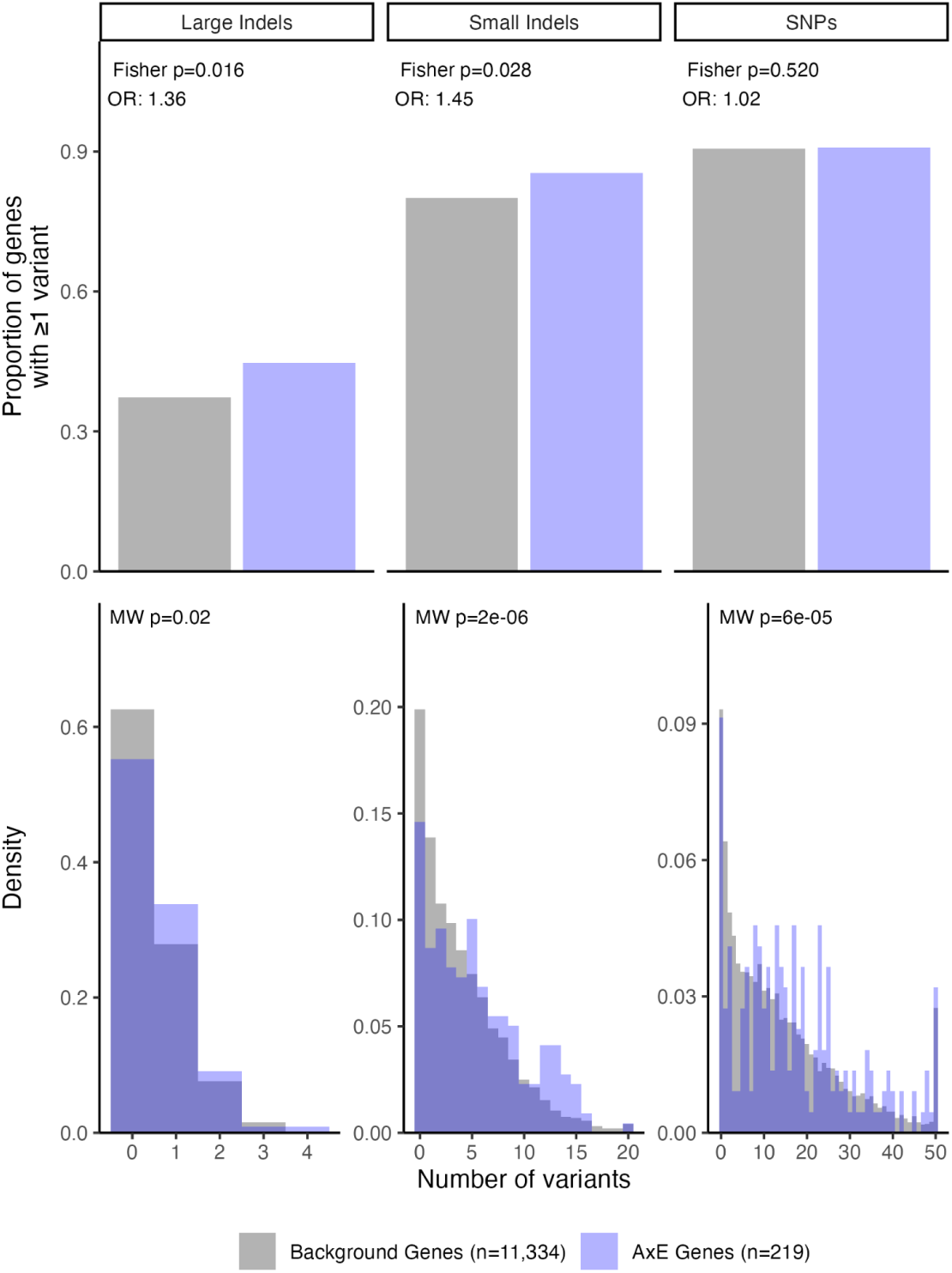
AxE genes (blue) are enriched for sequence variants compared to background genes (gray). We tested for enrichment of SNPs, small indels (<50bp), and large indels (≥50bp) in the promoters of genes (+/-500bp from the annotated transcription start site). Top row: one-sided Fisher’s Exact tests for presence/absence of each category of sequence variant in the promoter. Bottom row: one-sided Mann-Whitney U test for number of variants in the promoter.

### Genotype-specific DAP-seq peaks are enriched near AxE genes and overlap with small indels

To test whether GxE expression patterns are attributable to differential transcription factor binding between alleles, we used publicly available DAP-seq data for 198 TFs assayed in inbreds B73 and Mo17 (Galli, Chen, et al. 2025). DAP-seq is indicative of binding potential regardless of cell type or condition, and so represents a superset of possible binding sites(O’Malley et al. 2016). Though not all DAP-seq peaks are likely to be bound in a particular cell type or environment, the context-agnostic nature of DAP-seq is useful for accommodating the context mismatch between samples from this study and the samples used to generate the DAP-seq data.

Across all TFs, 102 AxE genes have at least one genotype-specific DAP-seq peak in their promoter, meaning the peak was observed in only one of the two parental inbred lines. This is an enrichment relative to background genes (Fisher’s test odds ratio = 1.47, p-value = 3e-3; Figure 3A; Supplementary File 4). AxE genes also tend to have more genotype-specific DAP-peaks than background genes (Mann-Whitney p=5e-3; Figure 3B). Among 115 individual TFs that had at least one genotype-specific peak in an AxE gene promoter, we found 91 of them, or 79%, are enriched in AxE genes (Fisher’s odds ratio > 1; Figure 3C; Supplementary File 5). These results show that AxE genes are more likely to have genotype-specific TF binding than background genes.

**Figure 3:**
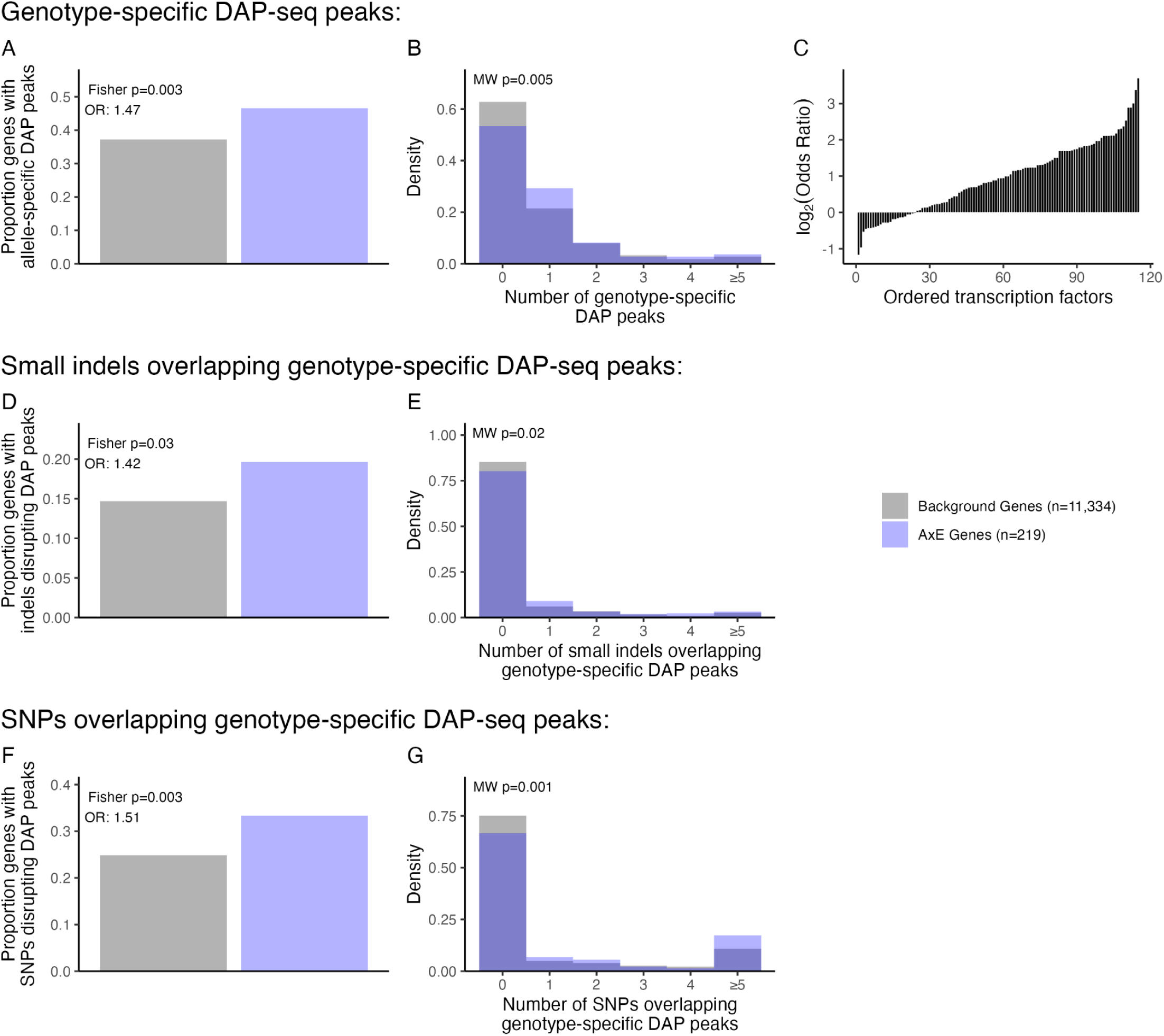
Genotype-specific DAP-seq peaks are more frequent, and more frequently disrupted, in AxE genes (blue). Relative to background (gray), AxE genes’ promoters (+/-500bp from TSS) are A) enriched for presence of one or more genotype-specific DAP-seq peaks and B) tend to have more genotype-specific DAP-seq peaks. C) When testing DAP-seq peaks for each transcription factor individually rather than in aggregate, the majority of transcription factors are enriched, rather than depleted, for genotype-specific binding sites in promoters of AxE genes (Fisher’s Odds Ratio > 1). Tests for whether small indels (<50bp in length) and SNPs overlap genotype-specific DAP-seq peaks show that AxE genes are D, F) enriched for one or more indels/SNPs overlapping genotype-specific DAP-seq peaks and E, G) tend to have more indels/SNPs disrupting genotype-specific DAP-seq peaks than background genes.

Because we hypothesized that sequence differences drive genotypic-specific binding and ultimately cause GxE for gene expression, we next tested whether small indels and SNPs overlap with genotype-specific DAP-seq peaks more often in AxE genes than in background genes. We found that AxE genes are more likely to have DAP-seq peaks that are overlapped by small indels (n=47 AxE genes, Fisher’s odds ratio: 1.42, p=3e-2; Figure 3D) and SNPs (Fisher’s odds ratio: 1.51, p=3e-3; Figure 3F), and also have greater number of DAP-seq peaks overlapped by small indels (Mann-Whitney p=2e-2; Figure 3E) and SNPs (Mann-Whitney p=1e-3; Figure 3G) (Supplementary File 4). Because AxE genes show enrichment under all three tests - for sequence variants, genotype-specific DAP-seq peaks, and their intersect - we consider this supportive of the hypothesis that cis-regulation of transcriptional GxE is in part driven by differential TF binding between alleles.

These tests are based on a simplified model: we assume a single TF independently occupying a single motif which, when disrupted, reduces the ability of that TF to bind. However, it is also possible that sequence variation impacts more complicated modes of TF binding that might not be revealed directly by DAP-seq and might involve sequence variation that is not directly under the DAP-seq peak, such as higher-order protein structures (e.g., TF dimers) and proximal-distal interactions(Deplancke et al. 2016). Studies in human cell lines have shown that only 12% of allele-specific TF binding can be explained by sequence variants that disrupt canonical motifs, but allele-specific TF binding is strongly associated with sequence variation within 50bp of the binding peak(Reddy et al. 2012). In our data, across all tested genes, we find a similarly strong enrichment of variants: 6.66 variants on average within 50bp of genotype-specific DAP-seq peaks, compared to 4.17 variants near shared DAP-seq peaks (Mann-Whitney p=2.2e-99). In AxE genes, we see a similar but statistically insignificant enrichment (7.03 variants versus 5.74 variants, Mann-Whitney p=0.52), presumably due to low power with only 248 genotype-specific DAP-seq peaks in promoter regions, or because sequence variants are already enriched in AxE genes’ promoters (Figure 2).

### Enrichment results are stronger in AxE genes than in genes showing allele effects

Enrichment of genotype-specific DAP-seq peaks, small indels, and overlap between the two could be due to sequence differences driving marginal allele effects in genes which happen to overlap with AxE genes, rather than driving AxE specifically. For example, genes showing genetic variation for expression levels are enriched for promoter SNPs(Grishkevich et al. 2012), and in some cases the underlying causes of GxE for gene expression may overlap with causes of genetic variation in gene expression(Grishkevich and Yanai 2013).

Because GxE genes often have some degree of marginal genotypic regulation and might be regulated by the same mechanisms as genes with genotypic variation, we wanted to test whether the enrichment observed in AxE genes is driven simply by allele effects. To do this, we identified genes with significant allele (A) effects but no significant AxE (n=1,546); with significant AxE effects but not significant A (n=136); and genes with both significant A and AxE effects (A+AxE; n=83). We then ran the tests described in previous sections on each of those sets and compared the results.

We observed that A genes generally are more enriched than AxE genes for variants in their promoters, a trend previously observed in *C. elegans*(Grishkevich et al. 2012), while A+AxE genes have an intermediate value for SNPs in promoters but a higher enrichment than either A or AxE genes for small indels (Figure 4; Supplementary File 6).

**Figure 4:**
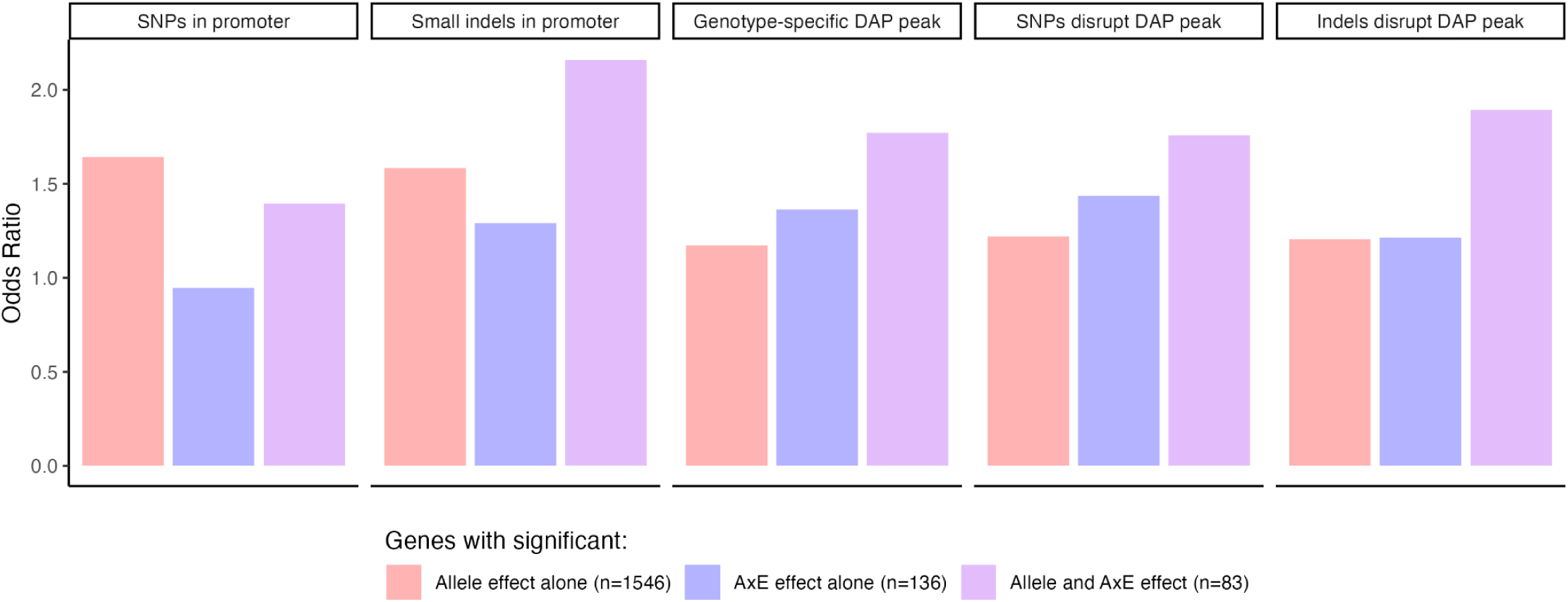
Enrichment of indels, SNPs, and DAP-seq peaks are greater in AxE genes than genes with only significant allele effects. Bars show Fisher’s test Odds Ratios, similar to tests in Figures 2 (top middle and top right panels), 3A, 3D, and 3F, for enrichment of small indels, SNPs, genotype-specific DAP-peaks, and indels/SNPs disrupting genotype-specific DAP peaks. We tested genes that show significant allele effects but no AxE (red), genes that show AxE effects but no allele effect (blue), and genes with significant allele and AxE effects (purple).

However, AxE genes show stronger enrichment than A genes for genotype-specific DAP-seq peaks and genotype-specific DAP-seq peaks disrupted by SNPs (Figure 4; Supplementary File 6). A+AxE genes had even higher odds ratios than A or AxE genes for genotype-specific DAP-seq peaks and genotype-specific DAP-seq peaks disrupted by indels or SNPs. We hypothesize that genes with both A and AxE effects are some of the most strongly regulated genes in these sets, driving their high enrichment for putative regulatory variation observed in this study.

### Sequence variation and DAP-seq features explain a modest amount of gene-proximal regulation of AxE

Finally, we used sequence variation and DAP-seq data to further test our hypothesis that sequence variation and TF binding differences contribute to transcriptional GxE. If so, localized promoter features should be able to predict which genes showed transcriptional AxE in this study. We also reasoned that the predictive ability of these models would give some indication of the degree to which sequence and TF binding drive AxE, versus other sources of gene regulation untested by this study.

Classification models using LASSO, Support Vector Machines, Random Forest, and Logistic Regression achieved modest overall predictive capacities, with ROC-AUC scores ranging between 0.54 and 0.60 (Supplementary Table 1). Although these models confirmed the presence of an underlying biological signal, they lacked consensus regarding feature importance; the LASSO framework identified B73-specific DAP-seq features as primary drivers, whereas other model architectures favored number of SNPs or Mo17-specific peaks, consistent with known genome-wide transcription factor binding site divergences between these two inbred lines(Galli, Chen, et al. 2025).

Given low agreement on feature importance and modest accuracy, we evaluated the standalone predictive power and univariate importance of individual features and identified eight features that exhibit minor but statistically significant signals for distinguishing AxE genes, with standalone ROC-AUC ranging from 0.551 to 0.591(Supplementary Table 2, Supplementary Figure 1). These eight features consolidate into three categories: total promoter SNP density, small indel frequency, and the fraction of genotype-specific DAP-seq peaks. This suggests that more polymorphism within a gene’s promoter region is a key indicator of AxE potential. Such a relationship aligns with foundational observations that extensive allelic cis-regulatory variation between B73 and Mo17 dictates downstream expression divergence(Stupar and Springer 2006).

While these results establish a statistical association between cis-regulatory variation and AxE classification, their standalone predictive power remains limited. Even when these features are combined in machine learning architectures, model performance maxes out at an ROC-AUC of 0.603 (Supplementary Table 1). These findings demonstrate that while promoter region sequence variation and TF binding potential establish a baseline for predicting AxE, improving predictive ability and extending it beyond hybrid genotypes will likely require expanding the modeling framework to incorporate trans-eQTL networks, epigenetic dynamics, or broader macromolecular interactions.

## Conclusions

In this study, we used allele-specific expression in B73xMo17 hybrids with uniform gene-distal regulatory background to focus on gene-proximal effects driving AxE for gene transcription. We found that compared to background genes and genes showing only genotype-specific variation, genes showing AxE interactions for transcript abundance have more small indels and SNPs in their promoters; more genotype-specific DAP-seq peaks; and that sequence variants disrupt the genotype-specific DAP-seq peaks more frequently. These results provide evidence supporting the hypothesis that sequence divergence in regulatory regions can drive GxE for gene expression. Though much of GxE for gene expression is regulated in trans, this study allows a novel, gene-proximal perspective on a mechanistic explanation of GxE in gene expression which can inform future work to model and engineer gene expression patterns across genetic backgrounds and environmental contexts.

## Materials & Methods

### Germplasm and sample collection

B73xMo17 hybrids were grown as hybrid check plots, randomly distributed throughout another experiment in replicated field trials in Clayton, NC and Columbia, MO in 2024. The NC experiment was planted April 16 and tissue for RNAseq was sampled on May 21, 36 days and 659.15 growing degree units (GDUs) after planting, between 10:08am and 10:46am (4.1-4.7hr after sunrise). The MO experiment was planted May 20 and sampled on June 13, 25 days and 493.05 GDUs after planting, between 9:11am and 12:15am (3.5-6.5hr after sunrise). A Midco Tissue Punch was used to collect one punch per leaf from five representative plants per plot into a 2mL tube containing 2 5/32” grinding beads. Samples were flash frozen in liquid nitrogen immediately after collection, and stored in a -80 freezer after.

### RNA extraction and sequencing

Prior to extraction, samples were ground using a Genogrinder 2010 Homogenizer at 1200 rpm for 20 seconds. RNA was extracted using the MagMAX™ Plant RNA Isolation Kit and the Kingfisher Flex. Library preparation and sequencing was performed by Novogene Corporation Inc. Briefly, messenger RNA was purified from total RNA using poly-T oligo-attached magnetic beads. After fragmentation, the first strand cDNA was synthesized using random hexamer primers followed by the second strand cDNA synthesis. Library end repair, A-tailing, adapter ligation, size selection, amplification, and purification were performed before 150bp paired-end sequencing on an Illumina NovaSeq X Plus.

### Allele specific RNAseq counts

A VCF file containing biallelic heterozygous SNPs for allele-specific read counting was generated following the workflow described in the AnchorWave GATK documentation (https://github.com/baoxingsong/AnchorWave/blob/master/doc/GATK.md). Briefly, the B73 v5 and Mo17-CAU-2 reference genomes were downloaded from MaizeGDB (https://download.maizegdb.org/), and the Mo17 genome was aligned to B73 using the “genoAli” function in AnchorWave v1.2.6(Song et al. 2022). The resulting AnchorWave MAF output was converted to GVCF format using the TASSEL MAFToGVCFPlugin(Bradbury et al. 2007) and processed using the GATK workflow to generate a final VCF file. This VCF file was further processed with TASSEL to generate the hybrid genotype file used for allele-specific counting.

RNA-seq reads were trimmed using fastp v1.3.3(Chen et al. 2018; Chen 2025), and read quality was assessed before and after trimming using FastQC v0.12.1 (https://www.bioinformatics.babraham.ac.uk/projects/fastqc/). Aggregated quality-control reports were generated using MultiQC v1.34(Ewels et al. 2016). We used the methods described in Hu et al. (2022)(Hu et al. 2022) to generate mapping-bias-corrected allele-specific counts for hybrid samples. Briefly, the B73 v5 reference genome and corresponding annotation were used to build a genome index using STAR v2.7.11b(Dobin et al. 2013). Trimmed reads from each sample were then aligned in two-pass mode and filtered to retain uniquely mapped reads with mapping quality ≥40. Allelic reads from mapped BAM files were counted at each heterozygous SNP site using ASEReadCounter from GATK v4.6.0.0(McKenna et al. 2010; Castel et al. 2015). To reduce the effects of low-depth sampling, false heterozygous calls caused by sequencing error, and reference-mapping bias, we retained only SNPs in each RNA-seq sample where both alleles were detected, total read depth was at least 10, and the absolute log2 ratio of reference to alternate allele counts was ≤2. The raw VCF was then subset to these positions to generate a sample-specific VCF. Reads were realigned with STAR using the same two-pass strategy but with the WASP filter enabled (‘--waspOutputMode SAMtag’) and the sample-specific VCF supplied via ‘--varVCFfilè, and only reads passing the WASP mapping-bias filter (‘vW:i:1’) were retained(Dobin et al. 2013; van de Geijn et al. 2015; Asiimwe and Alexander 2024). We extracted reads assigned to either B73 or Mo17 at all overlapping loci into separate BAM files and then counted the reads overlapping each gene feature in each BAM file using featureCounts v2.0.2(Liao et al. 2014). These gene counts represent the allelic expression of the B73 and Mo17 alleles of each gene, respectively.

### Identifying AxE genes

We tested for AxE as a change in the ratio of Mo17:B73 read counts at each gene, following the procedure outlined in (Love 2017). Read counts were modeled using a negative binomial distribution with DESeq2(Love et al. 2014) according to Equation 1:

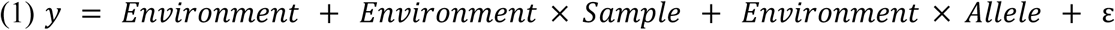

Where *Environment* is a fixed term for the location, Missouri or North Carolina; *Environment* × *Sample* is a fixed effect to account for variation among samples within each environment; *Environment* × *Allele* is a fixed effect to estimate Mo17:B73 ratios separately for each environment; and ε is an independent and identically distributed residual term. Size factors were uniformly set to 1, since the model is testing for differences of count ratios within each sample.

We tested for genes with significant AxE by contrasting the allelic ratios in each environment. Genes with FDR p<0.1 were considered significant AxE genes. Because there is not an explicit test for genetic (A) effects, or persistent differences between alleles, in Equation 1, we identified significant A genes as those with 1) significant environment-specific allelic ratios (FDR p<0.1) in both environments and 2) allelic ratios favoring the same allele in both environments (ie, either B73 or Mo17 higher in both).

The set of background genes against which AxE and A genes were compared was defined as all genes that 1) were tested in the DESeq2 model (assigned a p-value), and 2) had >10 counts in >5 columns, where a column represents one allele in one sample. The intent of this filtering was to ensure that background genes were sufficiently transcribed and to avoid including pseudogenes or other annotation errors that might inflate the apparent differences between A/AxE genes and background genes.

### Testing for overlap with phenotypic GxE

We used previously published phenotypic data from the intermated B73xMo17 population (IBM), consisting of 200 recombinant inbred lines (RILs) grown in up to 10 environments (number of environments varied by trait) and phenotype for 19 distinct traits(Hung et al. 2012). For genetic data, we used genotyping-by-sequencing (GBS)(Elshire et al. 2011) variant calls from the maize HapMap3(Bukowski et al. 2018). Genotype calls for IBM individuals were filtered to only sites that segregate between B73 and Mo17, then RILs with multiple replicates in the genotyping data were merged and RILs with <95% agreement rate between replicates were dropped from analysis. We kept individuals with <80% missing data and variants with <80% missing data and allele frequencies between 0.25 and 0.75. Variant positions on B73v4(Jiao et al. 2017) were uplifted to v5(Hufford et al. 2021) using the chain file available at https://download.maizegdb.org/Zm-B73-REFERENCE-NAM-5.0/chain_files/B73_RefGen_v4_to_Zm-B73-REFERENCE-NAM-5.0.chain. Genotypes were imputed using R/qtl2(Broman et al. 2019), and individuals with more recombinations than the 90th percentile of the population were removed. We identified 15,092 recombination bins across the remaining 208 individuals and created a samples x bins rectangular matrix to use for testing for GxE QTL.

To test for GxE QTL, we first removed block effects from the reported phenotypic data and used those block-adjusted phenotypes for mapping. We fit a null model to each trait (Equation 2) that accounts for polygenic line effects and environmental effects, where *EntryID* is a fixed effect for a particular RIL and *Environment* is a fixed effect for a given location/year combination. The error term ε is independent and identically distributed. We compared the null model to a model that included a GxE term (Equation 3), where terms are the same as in Equation 2 but with the addition of *Marker* × *Environment*, a fixed effect testing for environment-specific effects of the B73 or Mo17 allele in each recombination bin.

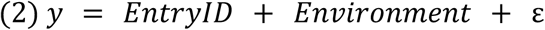

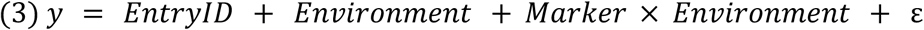

We scanned for GxE QTLs for each trait and estimated a combined p-value across all traits for each recombination bin using Stouffer’s method for combining p-values(Stouffer and Hovland 1949). Significant GxE QTL for individual traits as well as the GxE QTL identified by Stouffer’s method were tested by a Fisher’s exact test for enrichment of AxE genes identified from our gene expression data.

### Testing for enrichment of sequence variants and DAP-seq peaks

Structural variants between B73 and Mo17 were identified using the method described by(Galli, Chen, et al. 2025). The AnchorWave MAF alignment output was converted to PAF format using wgatools v1.1.0(Wei et al. 2025). The resulting PAF file was used as input for SyRI v1.7.1(Goel et al. 2019), and the resulting VCF file was parsed into SNPs, small indels (<50 bp), and large indels (≥50 bp).

To test enrichment of variants and DAP-seq peaks in gene promoters, we first defined promoters as ranging from 500bp upstream to 500bp downstream of the annotated transcription start site of each gene in the Zm-B73-REFERENCE-NAM-5.0_Zm00001eb.1 genome annotation of B73(Hufford et al. 2021). We counted the number of each variant type previously described and DAP-seq peaks from the file “normalized_specific_and_shared_peaks.zip”(Galli, Chen, et al. 2025; Galli, Huang, et al. 2025) overlapping promoters. To test for genotype-specific DAP-seq peaks disrupted by small indels, we counted the number of genotype-specific DAP-seq peaks that overlap at least one small indel in each promoter. Test for enrichment based on presence/absence was done with a one-sided Fisher’s exact test, and testing for differences in the number of features was done with a one-sided Mann-Whitney test.

### Prediction of GxE status from sequence features

Datasets containing transcription factor binding profiles (DAP-seq) and sequence variation features were compiled to predict binary AxE gene status using the tidymodels (https://www.tidymodels.org) ecosystem in R. Categorical predictors were dummy-coded, continuous numeric features were center-and-scale normalized, and highly collinear variables were removed using a pairwise correlation threshold of |r|>0.85(Vatcheva et al. 2016). To address baseline class imbalance, training data partitions were downsampled prior to model fitting(He and Garcia 2009; Kuhn and Johnson 2013).

Global classification performance was evaluated using stratified 5-fold cross-validation across four optimized architectures:

● *LASSO Regression:* Implemented via glmnet with a pure L1 penalty (mixture = 1.0) tuned across a 50-level regular lambda grid(Friedman et al. 2010).
● *Support Vector Machine (SVM):* Implemented via kernlab using a Radial Basis Function (RBF) kernel tuned across cost and sigma grids(Karatzoglou et al. 2004).
● *Random Forest:* Implemented via ranger with 500 decision trees using permutation-based variable importance calculations(Wright and Ziegler 2017).
● *Logistic Regression:* Configured as an unpenalized generalized linear model via core glm(R Core Team 2018).

Hyperparameters were optimized to maximize the Receiver Operating Characteristic Area Under the Curve (ROC-AUC) using the select_best criterion.

Univariate screening was concurrently executed to evaluate the standalone predictive power and information-theoretic value of each feature independently. Standalone ROC-AUC scores were calculated using the pROC package(Robin et al. 2011), where nominal predictors were pre-processed via binomial generalized linear models to generate conditional probability vectors. Non-linear dependencies were evaluated by calculating Shannon-based Mutual Information (MI) scores across empirical joint contingency matrices; continuous features were dynamically discretized into a maximum of ten equal-width bins prior to MI calculation(Shannon 1948). Features were ranked hierarchically by absolute divergence from random chance (|ROC-AUC=0.50|) and descending MI scores.

To robustly isolate cis-regulatory predictors, wrapper-based feature selection was implemented using the Boruta package(Kursa and Rudnicki 2010). The algorithm was executed over 250 iterations using a Random Forest classifier engine (ranger), comparing real feature importance against permuted shadow attributes. Statistical thresholds were strictly controlled using a Bonferroni multi-comparison adjustment(Dunn 1961). Features were definitively classified as confirmed, tentative, or rejected based on their mean importance Z-scores relative to the maximal shadow feature baseline.

### Use of generative artificial intelligence tools

Generative artificial intelligence (Claude Sonnet v4.6 and Opus 4.8; ChatGPT-5.5; Gemini v1.5 Pro) was used to assist with scripting, troubleshooting, and documenting code used for this project. Additionally, AI was used to edit writing for grammar, clarity, and flow. Outputs were reviewed for integrity by human authors of this publication.

## Data Availability Statement

Analyses from this study can be reproduced using code at https://github.com/gagelab/bmo_ase_gxe. Sequencing data have been deposited at NCBI SRA under <INPREPARATION>.

## Acknowledgements

Thanks to the staff at Central Crops Research Station and Genetics Farm for facilitating field experiments. Thanks to the North Carolina State University High Performance Computing Services Core Facility (RRID:SCR_022168) for supporting computing, and to the North Carolina State University Genomic Sciences Laboratory Core Facility (RRID:SCR_026944) for facilitating RNA extractions.

## References

1. Albert E, Duboscq R, Latreille M, Santoni S, Beukers M, Bouchet J-P, Bitton F, Gricourt J, Poncet C, Gautier V, et al. 2018. Allele-specific expression and genetic determinants of transcriptomic variations in response to mild water deficit in tomato. Plant J. 96(3):635–650.

2. Asiimwe R, Alexander D. 2024. STAR+WASP reduces reference bias in the allele-specific mapping of RNA-seq reads. bioRxivorg. doi:10.1101/2024.01.21.576391. http://biorxiv.org/lookup/doi/10.1101/2024.01.21.576391.

3. Ballinger MA, Mack KL, Durkin SM, Riddell EA, Nachman MW. 2023. Environmentally robust cis-regulatory changes underlie rapid climatic adaptation. Proc Natl Acad Sci U S A. 120(39):e2214614120.

4. Barreiro LB, Tailleux L, Pai AA, Gicquel B, Marioni JC, Gilad Y. 2012. Deciphering the genetic architecture of variation in the immune response to Mycobacterium tuberculosis infection. Proc Natl Acad Sci U S A. 109(4):1204–1209.

5. Boye C, Nirmalan S, Ranjbaran A, Luca F. 2024. Genotype × environment interactions in gene regulation and complex traits. Nat Genet. 56(6):1057–1068.

6. Bradbury PJ, Zhang Z, Kroon DE, Casstevens TM, Ramdoss Y, Buckler ES. 2007. TASSEL: software for association mapping of complex traits in diverse samples. Bioinformatics. 23(19):2633–2635.

7. Broman KW, Gatti DM, Simecek P, Furlotte NA, Prins P, Sen Ś, Yandell BS, Churchill GA. 2019. R/qtl2: Software for mapping quantitative trait loci with high-dimensional data and multiparent populations. Genetics. 211(2):495–502.

8. Bukowski R, Guo X, Lu Y, Zou C, He B, Rong Z, Wang B, Xu D, Yang B, Xie C, et al. 2018. Construction of the third-generation Zea mays haplotype map. Gigascience. 7(4):1–12.

9. Castel SE, Levy-Moonshine A, Mohammadi P, Banks E, Lappalainen T. 2015. Tools and best practices for data processing in allelic expression analysis. Genome Biol. 16(1):195.

10. Chen Q, Liu Z, Wang B, Wang X, Lai J, Tian F. 2015. Transcriptome sequencing reveals the roles of transcription factors in modulating genotype by nitrogen interaction in maize. Plant Cell Rep. 34(10):1761–1771.

11. Chen S. 2025. fastp 1.0: An ultra-fast all-round tool for FASTQ data quality control and preprocessing. Imeta. 4(5):e70078.

12. Chen S, Zhou Y, Chen Y, Gu J. 2018. fastp: an ultra-fast all-in-one FASTQ preprocessor. Bioinformatics. 34(17):i884–i890.

13. Cowles CR, Hirschhorn JN, Altshuler D, Lander ES. 2002. Detection of regulatory variation in mouse genes. Nat Genet. 32(3):432–437.

14. Cubillos FA, Stegle O, Grondin C, Canut M, Tisné S, Gy I, Loudet O. 2014. Extensive cis-regulatory variation robust to environmental perturbation in Arabidopsis. Plant Cell. 26(11):4298–4310.

15. Deplancke B, Alpern D, Gardeux V. 2016. The genetics of transcription factor DNA binding variation. Cell. 166(3):538–554.

16. Dobin A, Davis CA, Schlesinger F, Drenkow J, Zaleski C, Jha S, Batut P, Chaisson M, Gingeras TR. 2013. STAR: ultrafast universal RNA-seq aligner. Bioinformatics. 29(1):15–21.

17. Dunn OJ. 1961. Multiple comparisons among means. J Am Stat Assoc. 56(293):52–64.

18. Elshire RJ, Glaubitz JC, Sun Q, Poland JA, Kawamoto K, Buckler ES, Mitchell SE. 2011. A robust, simple genotyping-by-sequencing (GBS) approach for high diversity species. PLoS One. 6(5):e19379.

19. Ewels P, Magnusson M, Lundin S, Käller M. 2016. MultiQC: summarize analysis results for multiple tools and samples in a single report. Bioinformatics. 32(19):3047–3048.

20. Fairfax BP, Humburg P, Makino S, Naranbhai V, Wong D, Lau E, Jostins L, Plant K, Andrews R, McGee C, et al. 2014. Innate immune activity conditions the effect of regulatory variants upon monocyte gene expression. Science. 343(6175):1246949.

21. Friedman J, Hastie T, Tibshirani R. 2010. Regularization paths for generalized linear models via coordinate descent. J Stat Softw. 33(1):1–22.

22. Galli M, Chen Z, Ghandour T, Chaudhry A, Gregory J, Feng F, Li M, Schleif N, Zhang X, Dong Y, et al. 2025. Transcription factor binding divergence drives transcriptional and phenotypic variation in maize. Nat Plants. 11(6):1205–1219.

23. Galli M, Huang S-SC, Gallavotti A. 2025. Data from: Transcription factor binding divergence drives transcriptional and phenotypic variation in maize. doi:10.5281/ZENODO.14991915. 10.5281/ZENODO.14991915.

24. van de Geijn B, McVicker G, Gilad Y, Pritchard JK. 2015. WASP: allele-specific software for robust molecular quantitative trait locus discovery. Nat Methods. 12(11):1061–1063.

25. Goel M, Sun H, Jiao W-B, Schneeberger K. 2019. SyRI: finding genomic rearrangements and local sequence differences from whole-genome assemblies. Genome Biol. 20(1):277.

26. Grishkevich V, Ben-Elazar S, Hashimshony T, Schott DH, Hunter CP, Yanai I. 2012. A genomic bias for genotype–environment interactions in C. elegans. Mol Syst Biol. 8(1):587.

27. Grishkevich V, Yanai I. 2013. The genomic determinants of genotype × environment interactions in gene expression. Trends Genet. 29(8):479–487.

28. He H, Garcia EA. 2009. Learning from imbalanced data. IEEE Trans Knowl Data Eng. 21(9):1263–1284.

29. Huang W, Carbone MA, Lyman RF, Anholt RRH. 2020. Genotype by environment interaction for gene expression in Drosophila melanogaster. Nature Communications. doi:10.1038/s41467-020-19131-y. https://www.nature.com/articles/s41467-020-19131-y.

30. Hufford MB, Seetharam AS, Woodhouse MR, Chougule KM, Ou S, Liu J, Ricci WA, Guo T, Olson A, Qiu Y, et al. 2021. De novo assembly, annotation, and comparative analysis of 26 diverse maize genomes. Science. 373(6555):655–662.

31. Hu H, Crow T, Nojoomi S, Schulz AJ, Estévez-Palmas JM, Hufford MB, Flint-Garcia S, Sawers R, Rellán-Álvarez R, Ross-Ibarra J, et al. 2022. Allele-specific expression reveals multiple paths to highland adaptation in maize. Mol Biol Evol. 39(11):msac239.

32. Hung H-Y, Browne C, Guill K, Coles N, Eller M, Garcia A, Lepak N, Melia-Hancock S, Oropeza-Rosas M, Salvo S, et al. 2012. The relationship between parental genetic or phenotypic divergence and progeny variation in the maize nested association mapping population. Heredity. 108(5):490–499.

33. Jakobson CM, Jarosz DF. 2019. Molecular origins of complex heritability in natural genotype-to-phenotype relationships. Cell Syst. 8(5):363–379.e3.

34. Jiao Y, Peluso P, Shi J, Liang T, Stitzer MC, Wang B, Campbell MS, Stein JC, Wei X, Chin C-S, et al. 2017. Improved maize reference genome with single-molecule technologies. Nature. 546(7659):524–527.

35. Karatzoglou A, Smola A, Hornik K, Zeileis A. 2004. kernlab-AnS4Package for Kernel Methods inR. J Stat Softw. 11(9):1–20.

36. van Kleunen M, Fischer M. 2005. Constraints on the evolution of adaptive phenotypic plasticity in plants: Research review. New Phytol. 166(1):49–60.

37. Kuhn M, Johnson K. 2013. Applied predictive modeling. 1st ed. New York, NY: Springer.

38. Kursa MB, Rudnicki WR. 2010. Feature Selection with theBorutaPackage. J Stat Softw. 36(11):1–13.

39. Kusmec A, de Leon N, Schnable PS. 2018. Harnessing phenotypic plasticity to improve maize yields. Front Plant Sci. 9:1377.

40. Landry CR, Hartl DL, Ranz JM. 2007. Genome clashes in hybrids: insights from gene expression. Heredity (Edinb). 99(5):483–493.

41. Landry CR, Oh J, Hartl DL, Cavalieri D. 2006. Genome-wide scan reveals that genetic variation for transcriptional plasticity in yeast is biased towards multi-copy and dispensable genes. Gene. 366(2):343–351.

42. Lee MN, Ye C, Villani A-C, Raj T, Li W, Eisenhaure TM, Imboywa SH, Chipendo PI, Ran FA, Slowikowski K, et al. 2014. Common genetic variants modulate pathogen-sensing responses in human dendritic cells. Science. 343(6175):1246980.

43. Lee M, Sharopova N, Beavis WD, Grant D, Katt M, Blair D, Hallauer A. 2002. Expanding the genetic map of maize with the intermated B73 × Mo17 (IBM) population. Plant Mol Biol. 48(5-6):453–461.

44. Lensink M, Monroe G, Kliebenstein DJ. 2025. Trans-regulatory loci shape natural variation of gene expression plasticity in Arabidopsis. Genetics. 230(4):iyaf116.

45. Liao Y, Smyth GK, Shi W. 2014. featureCounts: an efficient general purpose program for assigning sequence reads to genomic features. Bioinformatics. 30(7):923–930.

46. Liu S, Li C, Wang H, Wang S, Yang S, Liu X, Yan J, Li B, Beatty M, Zastrow-Hayes G, et al. 2020. Mapping regulatory variants controlling gene expression in drought response and tolerance in maize. Genome Biol. 21(1):163.

47. Li Y, Alvarez OA, Gutteling EW, Tijsterman M, Fu J, Riksen JAG, Hazendonk E, Prins P, Plasterk RHA, Jansen RC, et al. 2006. Mapping determinants of gene expression plasticity by genetical genomics in C. elegans. PLoS Genet. 2(12):e222.

48. Lovell JT, Schwartz S, Lowry DB, Shakirov EV, Bonnette JE, Weng X, Wang M, Johnson J, Sreedasyam A, Plott C, et al. 2016. Drought responsive gene expression regulatory divergence between upland and lowland ecotypes of a perennial C4 grass. Genome Res. 26(4):510–518.

49. Love M. 2017 May 12. RPubs - Using RNA-seq DE methods to detect allele-specific expression. [accessed 2026 Apr 24]. https://rpubs.com/mikelove/ase.

50. Love MI, Huber W, Anders S. 2014. Moderated estimation of fold change and dispersion for RNA-seq data with DESeq2. Genome Biol. 15(12):550.

51. Mack KL, Landino NP, Tertyshnaia M, Longo TC, Vera SA, Crew LA, McDonald K, Phifer-Rixey M. 2025. Gene-by-environment interactions and adaptive body size variation in mice from the Americas. Mol Biol Evol. 42(4):msaf078.

52. McKenna A, Hanna M, Banks E, Sivachenko A, Cibulskis K, Kernytsky A, Garimella K, Altshuler D, Gabriel S, Daly M, et al. 2010. The Genome Analysis Toolkit: a MapReduce framework for analyzing next-generation DNA sequencing data. Genome Res. 20(9):1297–1303.

53. Mora-Poblete F, Maldonado C, Henrique L, Uhdre R, Scapim CA, Mangolim CA. 2023. Multi-trait and multi-environment genomic prediction for flowering traits in maize: a deep learning approach. Front Plant Sci. 14:1153040.

54. Moyerbrailean GA, Richards AL, Kurtz D, Kalita CA, Davis GO, Harvey CT, Alazizi A, Watza D, Sorokin Y, Hauff N, et al. 2016. High-throughput allele-specific expression across 250 environmental conditions. Genome Res. 26(12):1627–1638.

55. Napier JD, Heckman RW, Juenger TE. 2023. Gene-by-environment interactions in plants: Molecular mechanisms, environmental drivers, and adaptive plasticity. Plant Cell. 35(1):109–124.

56. Nica AC, Dermitzakis ET. 2013. Expression quantitative trait loci: present and future. Philos Trans R Soc Lond B Biol Sci. 368(1620):20120362.

57. O’Malley RC, Huang S-SC, Song L, Lewsey MG, Bartlett A, Nery JR, Galli M, Gallavotti A, Ecker JR. 2016. Cistrome and Epicistrome Features Shape the Regulatory DNA Landscape. Cell. 165(5):1280–1292.

58. Promislow D. 2005. A regulatory network analysis of phenotypic plasticity in yeast. Am Nat. 165(5):515–523.

59. R Core Team. 2018. R: A Language and Environment for Statistical Computing. https://www.R-project.org/.

60. Reddy TE, Gertz J, Pauli F, Kucera KS, Varley KE, Newberry KM, Marinov GK, Mortazavi A, Williams BA, Song L, et al. 2012. Effects of sequence variation on differential allelic transcription factor occupancy and gene expression. Genome Res. 22(5):860–869.

61. Roberts M, Josephs EB. 2023. Weaker selection on genes with treatment-specific expression consistent with a limit on plasticity evolution in Arabidopsis thaliana. Genetics. 224(2):iyad074.

62. Robin X, Turck N, Hainard A, Tiberti N, Lisacek F, Sanchez J-C, Müller M. 2011. pROC: an open-source package for R and S+ to analyze and compare ROC curves. BMC Bioinformatics. 12(1):77.

63. Rockman MV, Kruglyak L. 2006. Genetics of global gene expression. Nat Rev Genet. 7(11):862–872.

64. Rogers AR, Dunne JC, Romay C, Bohn M, Buckler ES, Ciampitti IA, Edwards J, Ertl D, Flint-Garcia S, Gore MA, et al. 2021. The importance of dominance and genotype-by-environment interactions on grain yield variation in a large-scale public cooperative maize experiment. G3. 11(2). doi:10.1093/g3journal/jkaa050. 10.1093/g3journal/jkaa050.

65. Schadt EE, Monks SA, Drake TA, Lusis AJ, Che N, Colinayo V, Ruff TG, Milligan SB, Lamb JR, Cavet G, et al. 2003. Genetics of gene expression surveyed in maize, mouse and man. Nature. 422(6929):297–302.

66. Shannon CE. 1948. A mathematical theory of communication. Bell Syst Tech J. 27(3):379–423.

67. Shao L, Xing F, Xu C, Zhang Q, Che J, Wang X, Song J, Li X, Xiao J, Chen L-L, et al. 2019. Patterns of genome-wide allele-specific expression in hybrid rice and the implications on the genetic basis of heterosis. Proc Natl Acad Sci U S A. 116(12):5653–5658.

68. Siddiq MA, Duveau F, Wittkopp PJ. 2024. Plasticity and environment-specific relationships between gene expression and fitness in Saccharomyces cerevisiae. Nat Ecol Evol. 8(12):2184–2194.

69. Signor SA, Nuzhdin SV. 2018. The Evolution of Gene Expression in cis and trans. Trends Genet. 34(7):532–544.

70. Smith EN, Kruglyak L. 2008. Gene–environment interaction in yeast gene expression. PLoS Biol. https://journals.plos.org/plosbiology/article?id=10.1371/journal.pbio.0060083.

71. Song B, Marco-Sola S, Moreto M, Johnson L, Buckler ES, Stitzer MC. 2022. AnchorWave: Sensitive alignment of genomes with high sequence diversity, extensive structural polymorphism, and whole-genome duplication. Proc Natl Acad Sci U S A. 119(1):e2113075119.

72. Springer NM, Stupar RM. 2007. Allele-specific expression patterns reveal biases and embryo-specific parent-of-origin effects in hybrid maize. Plant Cell. 19(8):2391–2402.

73. Stouffer SA, Hovland CI. 1949. Studies in Social Psychology in World War II.

74. Stupar RM, Springer NM. 2006. Cis-transcriptional variation in maize inbred lines B73 and Mo17 leads to additive expression patterns in the F1 hybrid. Genetics. 173(4):2199–2210.

75. Teressa T, Semahegn Z, Bejiga T. 2021. Multi environments and genetic-environmental interaction (GxE) in plant breeding and its challenges: a review article. International Journal of Research Studies in Agricultural Sciences. 7(4):11–18.

76. Vatcheva KP, Lee M, McCormick JB, Rahbar MH. 2016. Multicollinearity in regression analyses conducted in epidemiologic studies. Epidemiology (Sunnyvale). 6(2):227.

77. Velotta JP, Cheviron ZA. 2018. Remodeling ancestral phenotypic plasticity in local adaptation: A new framework to explore the role of genetic compensation in the evolution of homeostasis. Integr Comp Biol. 58(6):1098–1110.

78. Waters AJ, Makarevitch I, Noshay J, Burghardt LT, Hirsch CN, Hirsch CD, Springer NM. 2017. Natural variation for gene expression responses to abiotic stress in maize. Plant J. 89(4):706–717.

79. Wei W, Gui S, Yang J, Garrison E, Yan J, Liu H-J. 2025. Wgatools: An ultrafast toolkit for manipulating whole-genome alignments. Bioinformatics. 41(4):btaf132.

80. Wittkopp PJ, Haerum BK, Clark AG. 2004. Evolutionary changes in cis and trans gene regulation. Nature. 430(6995):85–88.

81. Wittkopp PJ, Kalay G. 2011. Cis-regulatory elements: molecular mechanisms and evolutionary processes underlying divergence. Nat Rev Genet. 13(1):59–69.

82. Wright MN, Ziegler A. 2017. Ranger: A fast implementation of random forests for high dimensional data in C++ and R. J Stat Softw. 77(1):1–17.

83. Yan H, Yuan W, Velculescu VE, Vogelstein B, Kinzler KW. 2002. Allelic variation in human gene expression. Science. 297(5584):1143.

84. Yun J, Burnett AC, Rogers A, Des Marais DL. 2025. Genotype by environment interactions in gene regulation underlie the response to soil drying in the model grass Brachypodium distachyon. Mol Biol Evol. 42(10):msaf218.

